# Learning in visual regions as support for the bias in future value-driven choice

**DOI:** 10.1101/523340

**Authors:** Sara Jahfari, Jan Theeuwes, Tomas Knapen

## Abstract

Reinforcement learning can bias decision-making towards the option with the highest expected outcome. Cognitive learning theories associate this bias with the constant tracking of stimulus values and the evaluation of choice outcomes in the striatum and prefrontal cortex. Decisions however first require processing of sensory input, and to-date, we know far less about the interplay between learning and perception. This fMRI study (N=43), relates visual BOLD responses to value-beliefs during choice, and, signed prediction errors after outcomes. To understand these relationships, which co-occurred in the striatum, we sought relevance by evaluating the prediction of future value-based decisions in a separate transfer phase where learning was already established. We decoded choice outcomes with a 70% accuracy with a supervised machine learning algorithm that was given trial-by-trial BOLD from visual regions alongside more traditional motor, prefrontal, and striatal regions. Importantly, this decoding of future value-driven choice outcomes again highligted an important role for visual activity. These results raise the intriguing possibility that the tracking of value in visual cortex is supportive for the striatal bias towards the more valued option in future choice.

---

In decision-making, our value beliefs bias future choices. This bias is shaped by the outcomes of similar decisions made in the past where the action, or stimulus chosen, becomes associated with a positive or negative outcome (‘value beliefs’). The evaluation of value after an outcome, or the comparison of value in decisions, is traditionally associated with activity in the prefrontal cortex and striatum (O’Doherty et al. 2004, 2017; Daw et al. 2006; Kahnt et al. 2009; Hare et al. 2011; Jocham et al. 2011; Klein et al. 2017).

To underset the bias in action selection midbrain dopamine neurons are thought to send a teaching signal towards the striatum and prefrontal cortex after an outcome (Montague et al. 1996; Schultz et al. 1997; Tobler et al. 2005). In the striatum, future actions are facilitated by bursts in dopamine after positive outcomes or discouraged by dopamine dips after negative outcomes. The dorsal and ventral parts of the striatum are known to receive differential, but also overlapping, inputs from midbrain neurons (O’Doherty et al. 2004; Atallah et al. 2007). Ventral and dorsal striatum have also been ascribed a differential role during learning by reinforcement learning theories. Here, the ventral parts of the striatum are involved with the prediction of future outcomes through the processing of prediction errors, whereas the dorsal striatum uses the same information to maintain action values as a way to bias future actions towards the most favored option (Joel et al. 2002; Kahnt et al. 2009; Collins and Frank 2014). Intriguingly, however, before many of these value-based computations can take place, stimuli first have to be parsed from the natural world, an environment where most reward predicting events are perceptually complex. This suggests that sensory processing might be an important integral part of optimized value-based decision-making.

Here, we investigate whether choice outcomes can modulate the early sensory processing of perceptually complex stimuli to help bias future decisions. Recent neurophysiological studies find visually responsive neurons in the tail of the caudate nucleus, which is part of the dorsal striatum (Kim and Hikosaka 2013; Hikosaka et al. 2014). These neurons encode and differentiate stable reward values of visual objects to facilitate eye movements towards the most valued target, while at the same time inhibiting a movement towards the lesser valued object (Kim et al. 2017). Critically, differential modulations are also observed in the primary visual cortex where stronger cortical responses are seen for objects with higher values (Serences 2008; Serences and Saproo 2010), which is consistent with the response of visual neurons in the caudate. As visual cortex is densely connected to the striatum (Fernandez-Ruiz et al. 2001; Kravitz et al. 2013), prioritized visual processing of high-value stimuli could aid the integration of information regarding the most-valued choice in the striatum (Lim et al. 2011, 2013; Jahfari et al. 2015; Jahfari and Theeuwes 2017). To understand these visual-striatal interactions, we focus on a more detailed parsing of the underlying computations.

Specifically, we explored two questions by reanalyzing fMRI data from a probabilistic reinforcement learning task using faces as visual stimuli (Jahfari et al. 2018) (Figure 1a). First, we focus on the interplay between learning and visual activity in the fusiform face area (FFA) and occipital cortex (OC). Here, with the use of a Bayesian hierarchical reinforcement learning model (Figure 1b) we outline how trial-by-trial estimates of action values (*Q*-value) and reward prediction errors (RPE) relate to the BOLD response of visual regions and the striatum (O’Doherty et al. 2007; Daw 2011) (Figure 1c). Second, we analyze data from a follow-up transfer phase, where the learning of value was already established. In our analysis, the importance of visual brain activity in the prediction, or decoding, of future value-based decisions is evaluated by using a supervised Random Forest (RF) machine learning algorithm (Breiman 2001, 2004). Specifically, transfer phase single-trial BOLD estimates from anatomically defined visual, prefrontal, and subcortical regions are combined by RF to predict, or decode, choice outcomes in a seperate validation set. We focus on classification accuracy, and the relative importance of each brain region in the correct classification of future value-based decisions.

**Figure 1:**
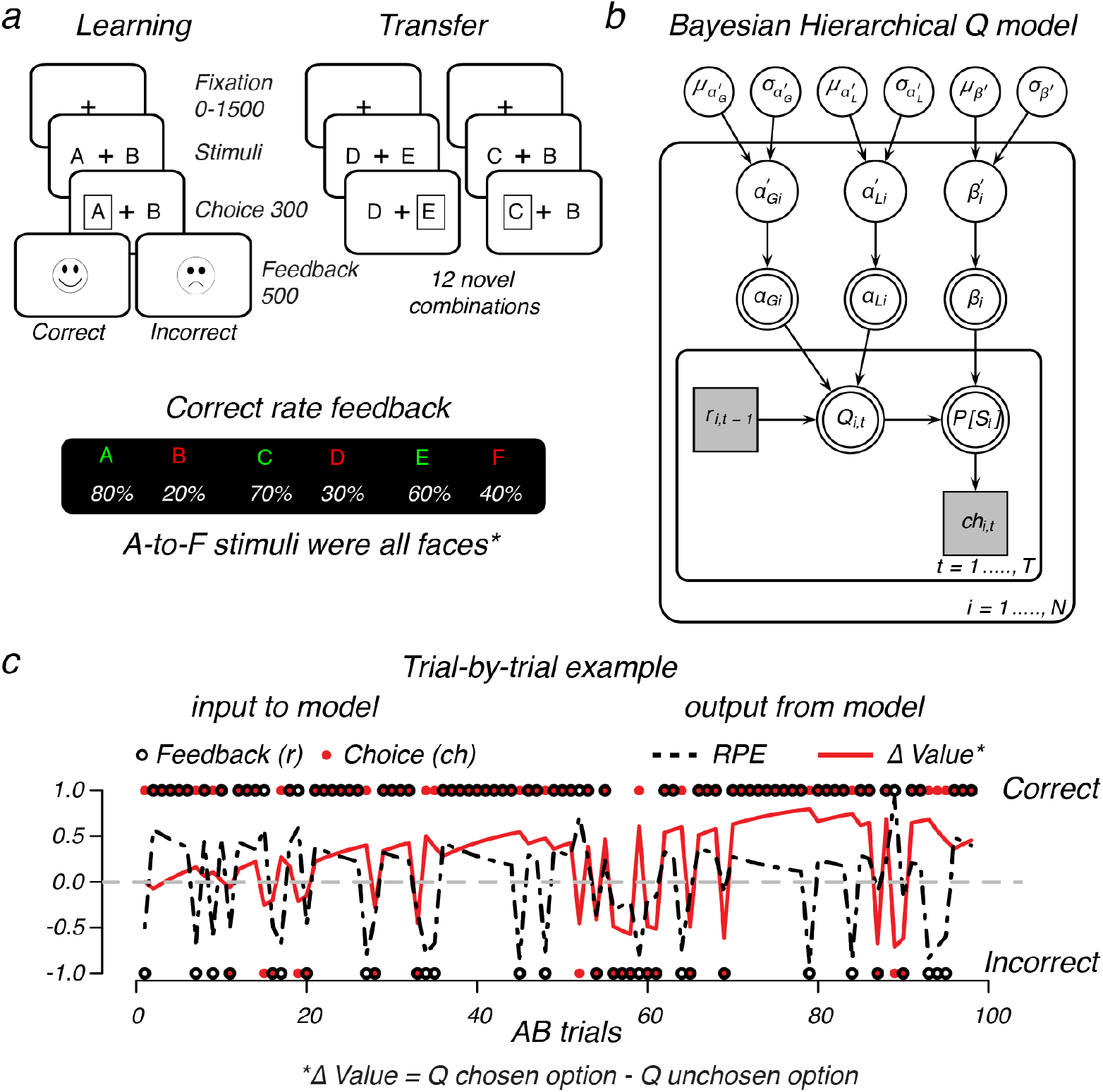
Design and Model. **a**) Reinforcement learning task using faces. During learning, two faces were presented on each trial, and participants learned to select the optimal face identity (A, C, E) through probabilistic feedback (% of correct is shown beneath each stimulus). The learning-phase contained three face pairs (AB, CD, ED) for which feedback was given. In a follow-up transfer phase these faces were rearranged into 12 novel combinations to asses learning. These trials were identical to learning trials, with the exception of feedback. *Example faces were removed for the publication on BioRxiv, for an impression see Jahfari et al. (2018), or the Radboud face database from where the faces were originally selected (http://www.socsci.ru.nl:8180/RaFD2/RaFD). **b**) Graphical *Q*-learning model with hierarchical Bayesian parameter estimation. The model consists of an outer subject (*i* = 1,. . . . ., *N*), and an inner trial plane (*t* = 1, …, *T*). Nodes represent variables of interest. Arrows are used to indicate dependencies between variables. Double borders indicate deterministic variables. Continuous variables are denoted with circular nodes, and discrete with square nodes. Observed variables are shaded in grey (see methods for details about the fitting procedure). **c**) Illustration of the observed trial-by-trial input (i.e., the choice made, and feedback received), and output (i.e., *Q* for the chosen and unchosen stimulus, ∆Value, and RPE) of the model given the estimated variability in learning rates from either positive (*α*_*Gi*_) or negative (*α*_*Li*_) feedback, and the tendency to exploit *β* higher values *i*.

## Materials and Methods

To understand how value learning relates to the activity pattern in perceptual regions we reanalyzed the behavioral and fMRI recordings of a recent study (Jahfari et al. 2018). In this study, BOLD signals were recorded while participants performed a reinforcement learning task using male or female faces, and a stop-signal task (which was discussed in Jahfari et al. (2018)). The fusiform face area (FFA) was localized using a separate experimental run.

### Participants

49 young adults (25 male; mean age = 22 years; range 19-29 years) participated in the study. All participants had normal or corrected-to-normal vision and provided written consent before the scanning session, in accordance with the declaration of Helsinki. The ethics committee of the University of Amsterdam approved the experiment, and all procedures were in accordance with relevant laws and institutional guidelines. In total, six participants were excluded from all analyses due to movement (2), incomplete sessions (3), or misunderstanding of task instructions (1). In total data from 43 participants was analyzed.

### Reinforcement learning task

Full details of the reinforcement learning task are provided in Jahfari et al. (2018). In brief, the task consisted of two phases (Figure 1a). In the first learning phase, three male or female face pairs (AB, CD, EF) were presented in a random order, and participants learned to select the most optimal face (A, C, E) in each pair solely through probabilistic feedback (‘correct’: happy smiley, ‘incorrect’: sad smiley). Choosing face-A lead to ‘correct’ on 80% of the trials, whereas a choice for face-B only lead to the feedback ‘correct’ for 20% of the trials. Other ratios for ‘correct’ were 70:30 (CD) and 60:40 (EF). Participants were not informed about the complementary relationship in pairs. All trials started with a jitter interval where only a white fixation cross was presented and had a duration of 0, 500, 1000 or 1500ms to obtain an interpolated temporal resolution of 500ms. Two faces were then shown left and right of the fixation-cross and remained on screen up to response, or trial end (4000ms). If a response was given on time, a white box surrounding the chosen face was then shown (300ms) and followed (interval 0-450ms) by feedback (500ms). Omissions were followed by the text ‘miss’ (2000ms). The transfer-phase contained the three face-pairs from the learning phase, and 12 novel combinations, in which participants had to select which item they thought had been more rewarding during learning. Transfer-phase trials were identical to the learning phase, with the exception that no feedback was provided. All trials had a fixed duration of 4000ms, where in addition to the jitter used at the beginning of each trial, null trials (4000ms) were randomly interspersed across the learning (60 trials; 20%) and transfer (72 trials; 20%) phase. Each face was presented equally often on the left or right side, and choices were indicated with the right-hand index (left) or middle (right) finger. Before the MRI session, participants performed a complete learning phase to familiarize with the task (300 trials with different faces). In the MRI scanner, participants performed two learning blocks of 150 trials each (300 trials total; equal numbers of AB, CD and EF), and three transfer phase blocks of 120 trials each (360 total; 24 presentations of each pair). All stimuli were presented on a black-projection screen that was viewed via a mirror-system attached to the MRI head coil.

### Reinforcement learning model

Trial-by-trial updating in value beliefs about the face selected in the learning phase, and reward prediction errors (signed expectancy violations) were estimated with a variant of the computational *Q*-learning algorithm (Watkins and Dayan 1992; Frank et al. 2007; Daw 2011) that is frequently used with this reinforcement learning task and contains two separate learning rate parameters for positive (*α*_*gain*_) and negative (*α*_*loss*_) reward prediction errors (Frank et al. 2007; Kahnt et al. 2009; Niv et al. 2012; Jahfari and Theeuwes 2017; Jahfari et al. 2018). *Q*-learning assumes participants to maintain reward expectations for each of the six (A-to-F) stimuli presented during the learning phase. The expected value (*Q*) for selecting a stimulus *i* (could be A-to-F) upon the next presentation is then updated as follows:

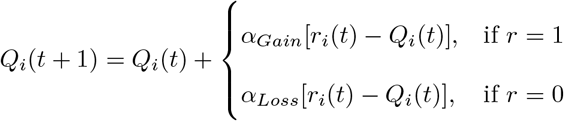

Where 0 ≤ *α*_*gain*_ or *α*_*loss*_ ≤ 1 represent learning rates, *t* is trial number, and *r* = 1 (positive feedback) or *r* = 0 (negative feedback). The probability of selecting one response over the other (i.e., A over B) is computed as:

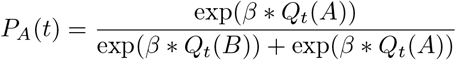

With 0 ≤ *β* ≤ 100 known as the inverse temperature.

### Bayesian hierarchical estimation procedure

To fit this *Q*-learning algorithm with two learning rate parameters we used Bayesian hierarchical estimation procedure. The full estimation procedure is explained in (Jahfari et al. 2018). To summarize, this implementation assumes that probit-transformed model parameters for each participant are drawn from a group-level normal distribution characterized by group level mean and standard deviation parameters: *z* ∼ *N* (*µ*_*z*_, *σ*_*z*_). A normal prior was assigned to group-level means *µ*_*z*_ ∼ *N* (0, 1), and a uniform prior to the group-level standard deviations *σ*_*z*_ ∼ *U* (1, 1.5). Model fits were implemented in Stan, where multiple chains were generated to ensure convergence.

### Image acquisition

The fMRI data for the Reinforcement learning task was acquired in a single scanning session with two learning and three transfer phase runs on a 3-T scanner (Philips Achieva TX, Andover, MA) using a 32-channel head coil. Each scanning run contained 340 functional *T*2^∗^-weighted echo-planar images for the learning phase, and 290 *T* 2^∗^-weighted echo planar images for the transfer phase (TR = 2000 ms; TE = 27.63 ms; FA = 76.1°; 3 mm slice thickness; 0.3 mm slice spacing; FOV = 240 × 121.8 × 240; 80 × 80 matrix; 37 slices, ascending slice order). After a short break of 10 minutes with no scanning, data collection was continued with a three-dimensional *T* 1 scan for registration purposes (repetition time [TR] = 8.5080 ms; echo time [TE] = 3.95ms; flip angle [FA] = 8°; 1 mm slice thickness; 0 mm slice spacing; field of view [FOV] = 240 × 220 × 188), the fMRI data collection using a stop signal task (described in Jahfari et al. (2018)), and a localizer task with faces, houses, objects, and scrambled scenes to identify FFA responsive regions on an individual level (317 *T* 2^∗^ weighted echo-planar images; TR = 1500 msec; TE = 27.6 msec; FA = 70°; 2.5 mm slice thickness; 0.25 mm slice spacing; FOV = 240 × 79.5 × 240; 96 × 96 matrix; 29 slices, ascending slice order). Here, participants viewed a series of houses, faces, objects as well as phase-scrambled scenes. To sustain attention during functional localization, subjects pressed a button when an image was directly repeated (12.5% likelihood).

### fMRI analysis learning phase

The interplay between learning and perceptual activity was examined by evaluating how trial-by-trial computations of value-beliefs, and reward prediction errors relate to BOLD responses in the occipital cortex (OC) and fusiform face area (FFA). To compare perceptual responses with the more traditional literature, we first show how value-beliefs and RPEs relate to the activity pattern of the dorsal (i.e., caudate, or putamen) or ventral (i.e., accumbens) parts of the striatum. Regions of interest (ROI) templates were defined using anatomical atlases available in FSL, or the localizer task for FFA. For this purpose, the localizer scans were preprocessed using motion correction, slice-time correction, and pre-whitening (Woolrich et al. 2001). For each subject, a GLM was fitted with the following EVs: for FFA, faces *>* (houses and objects), for parahippocampal place area (PPA), houses *>* (faces and objects) and for lateral occipital complex (LOC), intact scenes *>* scrambled scenes. Higher-level analysis was performed using FLAME Stage 1 and Stage 2 with automatic outlier detection (Beckmann et al. 2003). For the whole-brain analysis Z (Gaussianized T/F) statistic images were thresholded using clusters determined by *z >* 2.3 and *p* < .05 (GRFT) to define a group-level binary FFA region. Templates used for the caudate [center of gravity (cog): (-) 13, 10, 10], putamen [cog: (-) 25, 1, 1], and nucleus accumbens [cog: (-)19, 12, −7] were based on binary masks. Because participants were asked to differentiate faces, for each participant, we multiplied the binary templates of OC [cog: 1, −83, 5], FFA [cog: 23, −48, −18] with the individual t-stats from the localizer task contrast faces *>* (houses and objects). All anatomical masks, and the localizer group-level FFA mask can be downloaded from github (see acknowledgements).

### Deconvolution analysis learning phase

To more precisely examine the time course of activation in the striatal and perceptual regions, we performed finite impulse response estimation (FIR) on the BOLD signals. After motion correction, temporal filtering (3rd order savitzky-golay filter with window of 120 s) and percent signal change conversion, data from each region was averaged across voxels while weighting voxels according to ROI probability masks, and upsampled from 0.5 to 3 Hz. This allows the FIR fitting procedure to capitalize on the random timings (relative to TR onset) of the stimulus presentation and feedback events in the experiment. Separate response time courses were simultaneously estimated triggered on two separate events: stimulus onset, feedback onset. FIR time courses for all trial types were estimated simultaneously using a penalized (ridge) least-squares fit, as implemented in the FIRDeconvolution package (Knapen and Gee 2016), and the appropriate penalization parameter was estimated using cross-validation. For stimulus onset events (i.e., onset presentation of face pairs) response time courses were fit separately for the AB, CD and EF pairs, while also estimating the time courses of signal covariation with chosen and unchosen value for these pairs. For these events, our analysis corrected for the duration of the decision process. For the feedback events, the co-variation response time course with signed and unsigned prediction errors were estimated. These signal response time courses were analysed using across-subjects GLMs at each time-point using the statsmodels package (Seabold and Perktold 2010). The *α* value for the contributions of *Q* or RPE was set to 0.0125 (i.e. a Bonferroni corrected value of 0.05 given the interval of interest between 0 and 8 s).

### Random Forest classification

To specify the relevance of perceptual regions in the resolve of future value-driven choices a random forest (RF) classifier was used (Breiman 2001, 2004). The RF classifier relies on an ensemble of decision trees as base learners, where the final prediction (e.g., for a given trial is the choice going to be correct/optimal? or incorrect/suboptimal? given past learning) is obtained by a majority vote that combines the prediction of all decision trees. To achieve controlled variation, each decision tree is trained on a random subset of the variables (i.e. regions of interest chosen), and a bootstrapped sample of data points (i.e. trials or rows of the matrix in Figure 2c). In the construction of each tree about 1/3 of all trials is left out - termed as the “out-of-bag” sample – and later used to see how well each tree preforms on unseen data in the training set. Because in RF each tree is built from a different sample of the original data each observation is “out-of-bag” (OOB) for some of the trees. As such, each OOB sample is offered to all trees where the sample was not used for construction, and the average vote across those trees is taken as the classification outcome. The proportion of times that the classification outcome is not equal to the actual choice is averaged over all cases and represents the RF OOB error estimate. In other words, the generalized error for predictions is calculated by aggregating the prediction for every out-of-bag sample across all trees. In the results section, the OOB errors obtained from RF during training were well matched with the classification accuracy seen for the validation set given only the ‘good learners’ (OOB=30%, RF error validation set= 31%) or all participants (OOB= 33%, RF error validation set= 35%). An important feature of the RF classification method is the ease to measure the relative importance of each variable (i.e., region), in the overall predictive performance. That is, it allows for the ranking of all regions evaluated in the prediction of future value-based decisions.

**Figure 2:**
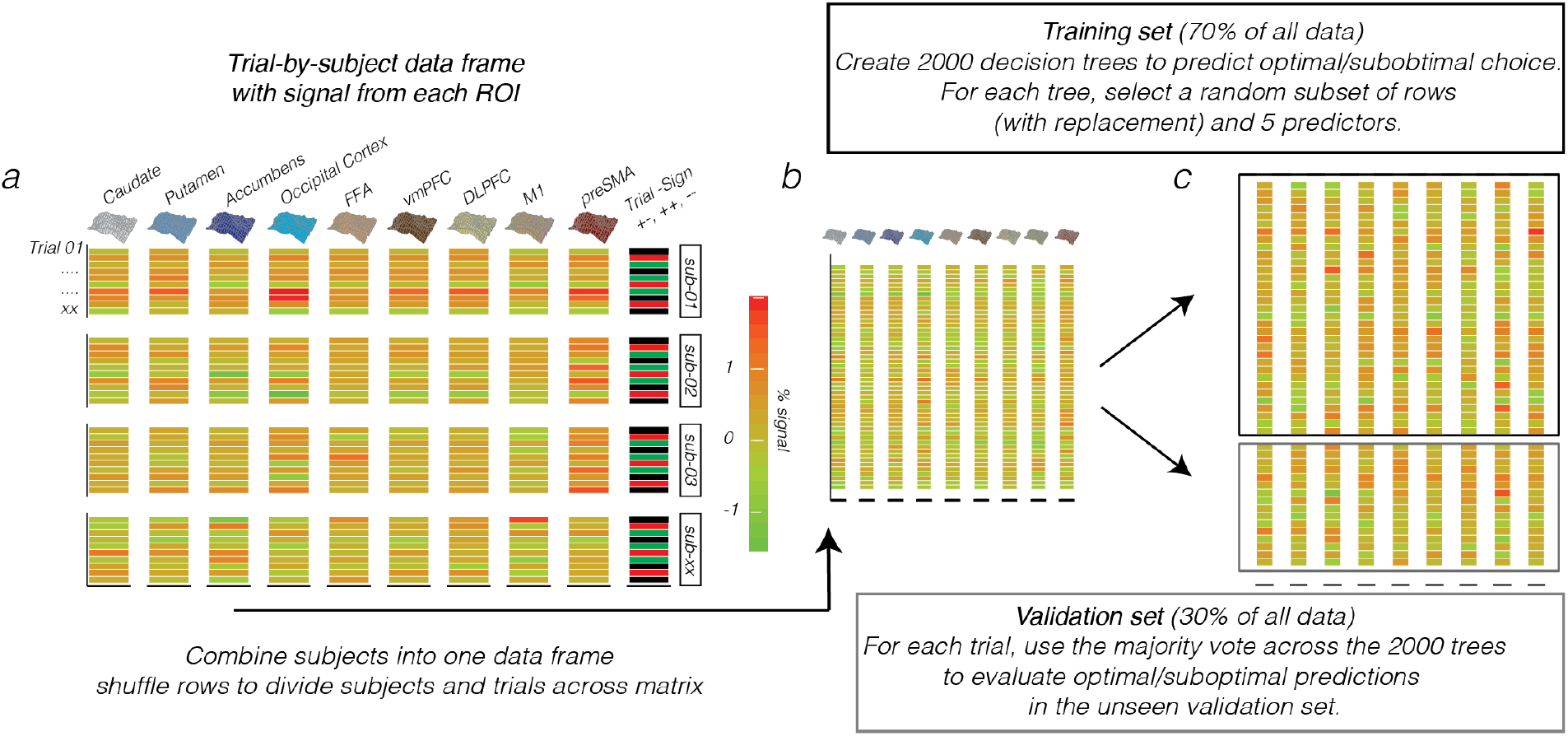
Random Forest input and data-structure. (**a**) Trial-by-subject data matrix with the % signal change drawn for each choice trial in the transfer-phase (rows) from 9 a-priori defined regions of interest (columns). In addition to the ROI data, the matrix contained a column with the identity of participants (sub-01, etc) and Trial Sign, which specified a choice between two positives (+/+; AC, AE, CE), negatives (−/−, BD, BF, DF), or between a negative and positive option (+/−, e.g., AD, CF, etc) given the feedback scheme in the learning-phase. (**b**) The individual subject data frames were then combined into one matrix, in which the rows were subsequently shuffled to randomly distribute trials and subjects across the rows. (**c**) This matrix was then divided into a training set (2/3 of the data) for the creation of 2000 decision trees of which the majority vote on each trial is then used to evaluate the predictive accuracy of optimal/suboptimal choices in a separate validation set (1/3 of the data).

### ROI selection and Random Forest procedure

This study used the ‘Breiman and Cutler’s Random Forests for Classification and Regression’ package in R, termed randomForest (randomForest_4.6-14). RF evaluations relied on the fMRI data recorded during the transfer phase, in a set of 9 regions of interest (ROIs). These ROIs included all templates from the learning phase (i.e., caudate, putamen, accumbens, OC, and FFA), as well as, the ventromedial prefrontal cortex (vmPFC), dorsolateral prefrontal cortex (DLPFC), pre-supplementary motor area (preSMA), and the primary motor cortex (M1). The selection of these additional anatomical templates was inspired by our previous analysis of this data with those templates focusing on networks (Pircalabelu et al. 2015; Schmittmann et al. 2015; Jahfari et al. 2018). Specifically, the DLPFC template was obtained from an earlier study, linking especially the posterior part to action execution (Cieslik et al. 2012). The preSMA, vmPFC, and M1 mask were created from cortical atalases available in FSL. Please notice that we used the same anatomical ROIs for both the model-based deconvolution analysis (Figure 4&5) and the decoding analysis (Figure 2&6).From each ROI a single parameter estimate (averaged normalized *β* estimate across voxels in each ROI) was obtained per trial, per subject. All, pre-processing steps to obtain single-trial images are described in Jahfari et al. (2018). Single-trial activity estimates were used as input variables in RF to predict choice outcomes (optimal/sub-optimal) in the transfer phase. Here, participants choose the best/optimal option based on values learned during the learning phase. We defined optimal choices as correct (i.e, when participants choose the option with the higher value), and sub-optimal choices as incorrect. Misses were excluded from RF evaluations.

By design, the transfer-phase contained 360 trials including 15 different pairs (12 novel), where each pair was presented 24 times with the higher value presented left in 12 of the 24 presentations, and on the right for the other half. With so many subtle value differences across the options presented and only one BOLD estimate per trial/region the prediction of future choices is under powered (Figure 2a). Therefore, assuming that all participants come from the same population, a fixed effects approach was taken for evaluations with RF. Here, the trial∗region activity matrices for all participants were combined into one big data matrix (Figure 2b) and subsequently shuffled across the rows, so that both participants and trials were re-arranged in a random order across rows. Besides the single trial BOLD estimates from the 9 ROI’s, this shuffled matrix contained two additional columns, which specified subject_id (to which subject does each trial belong), and Trial Sign – i.e., is the choice between the two faces about two positive (+/+; AC, AE, CE), negative (−/−; BD, BF, DF), or a positive-negative (+/−; e.g. AD, CF etc.) associations given the task manipulation during learning. Subject_id was included to control for different BOLD fluctuations across participants, whereas Trial Sign was added because both BOLD and choice patterns differ across these options (please see Jahfari et al. (2018)). The shuffled fixed effect matrix was divided into a separate training (2/3 of whole matrix), and validation (1/3) set, to be used for RF evaluations (Figure 2c). Based on our previous connectivity work with this data (Jahfari et al. 2018), we were aware that many of our single-trial BOLD response were correlated accross time, which potenentially results from shared learning effects (Supplementary figure 4). With RF the problem of correlated features is minimized for predictions with variable selection - i.e., the random selection of a set of regions to use for each tree. With more variables selected, we get better splits in each tree but also highly correlated decisions trees across the forest, which in essence diminishes the forest effect. To find the best balance, this study optimized the number of variables to select with a tuning function using the OOB error estimate. Learning was based on the training set, using 2000 trees with the number of variables (regions) used by each tree optimized with the tuneRF function in R, and accordingly set to 5. For the construction of each tree about 1/3 of all trials is left out - termed as the out-of-bag sample – and later used to see how well each tree preforms on unseen data. The generalized error for predictions is calculated by aggregating the prediction for every out-of-bag sample across all trees. Besides this out-of-bag approximation we evaluated the predictive accuracy of the whole RF on the separate unseen validation-set. We further reasoned that RF predictions can result from alternative BOLD patterns such as the buildup of a motor response, the ease of face distinctions, or to us alternative functional fluctuations. Therefore, prior to the evaluation of region importance (or ranking), we preformed two control analysis ensuring that RF predictions are sensitive to the consistency of past learning, and the representation of ∆*V alue*. These are the evaluations comparing ‘good’ to ‘all’ learners, as well as, the relationship between ∆*V alue* and RF uncertainty. In addition, while potential confounds of colinearity on the RF ranking cannot be excluded, we tried to minimize this with the use of permutation importance. Here, by using the OOB samples the importance of each variable (region) is computed as the difference between the models baseline accuracy and the drop in overall accuracy caused by permuting that column (region). While being more slow, permutation importance is described as more robust in comparison to the default (gini) importance computation where only the uncertainty of predictions is evaluated (with no checks on accuracy fluctuations after region permutation). The single trial data used as input, the RF evaluation codes, and ROI templates can all be downloaded from the github link provided in acknowledgements.

## Results

### Model and Behavior

As shown in Figure 1a, in the reinforcement learning task participants learned to select among choices with different probabilities of reinforcement (i.e., AB 80:20, CD 70:30, and EF 60:40). A subsequent transfer phase, where feedback was omitted, required participants to select the optimal option among novel pair combinations of the faces that were used during the learning phase (Figure 1a). In the learning phase, subjects reliably learned to choose the most optimal face option in all pairs. For each pair the probability of choosing the better option was above chance (*p*’s *<* .001), and the effect of learning decreased from AB (80:20) and CD (70:30) to the most uncertain EF (60:40) pair (*F* (2, 84) = 13.74, *p* < .0001). At the end of learning, value beliefs differentiating the optimal (A, C, E) from the sub-optimal (B, D, F) action were very distinct for the AB and CD face pairs but decreased with uncertainty (*F* (2, 84) = 39.70*, p <* 0.0001, Figure 3a). Value beliefs were estimated using the individual subject parameters of the *Q*-learning model that best captured the observed data (Figure 3b-e; reproduced from Jahfari et al. (2018) to show performance).

**Figure 3:**
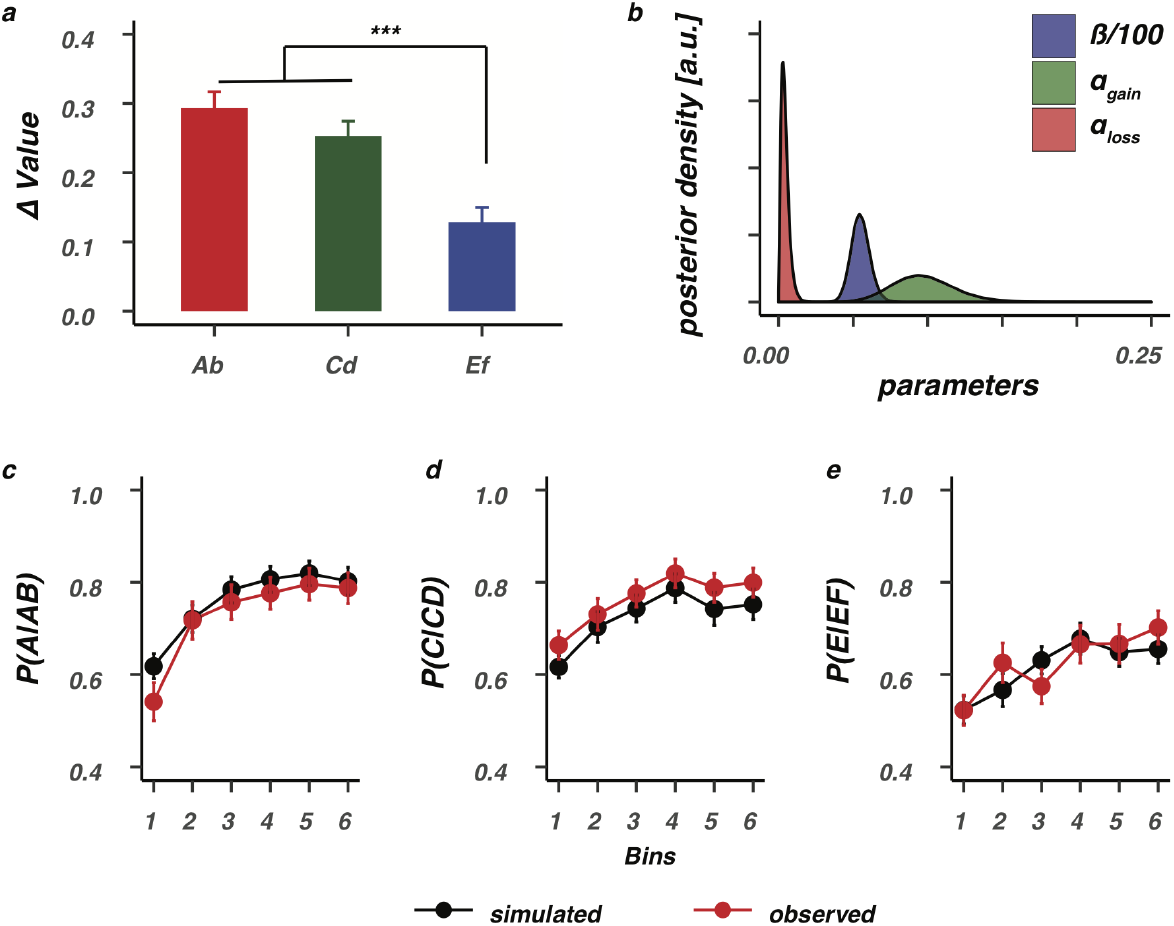
Value differentiation and model performance. (**a**) Value differentiation (∆Value) for the selection of the optimal (A,C,E) stimuli over the suboptimal (B,D,F) stimuli decreased as a function of feedback reliability, and was smallest for the most uncertain EF stimuli. *** = *p <* 0.0001, Bonferroni corrected. (**b**) Group-level posteriors for all *Q*-learning parameters. The bottom row shows model performance, where data was simulated with the estimated individual subject parameters and evaluated against the observed data for the AB (**c**), CD (**d**), or EF (**e**) pairs. Bins contain +/− 16 trials. Error bars represent standard error of the mean (SEM).

### BOLD is modulated by reliable value differences between faces in striatal and visual regions

For each pair of faces presented during the learning phase (AB, CD, EF) we asked how the BOLD signal time-course in striatal and visual regions relates to trial-by-trial value beliefs about the two faces presented as a choice. First, as a reference, we focused on the activity pattern of three striatal regions. Results showed BOLD responses in dorsal (caudate, putamen) but not ventral (accumbens) striatum to be differentially modulated by the estimated value beliefs of the chosen face (*Q*_*chosen*_), in comparison to value beliefs about the face that was not chosen (*Q*_*unchosen*_). Thus, BOLD responses in the dorsal striatum were modulated more strongly by value beliefs about the chosen stimulus (*Q*_*chosen*_; Figure 4a bottom row). Critically, this differential modulation was only observed with the presentation of AB faces where value differences were most distinct because of the reliable feedback scheme. Next, we evaluated the relationship between value and BOLD in the FFA, and OC. Again, only with the presentation of the AB face option, trial-by-trial BOLD fluctuations were differentially modulated by values of the chosen versus not chosen face option (Figure 4b bottom row). These evaluations highlight how the BOLD response in striatal and perceptual regions is especially sensitive to values of the (to-be) chosen stimulus when belief representations are stable and distinct.

**Figure 4:**
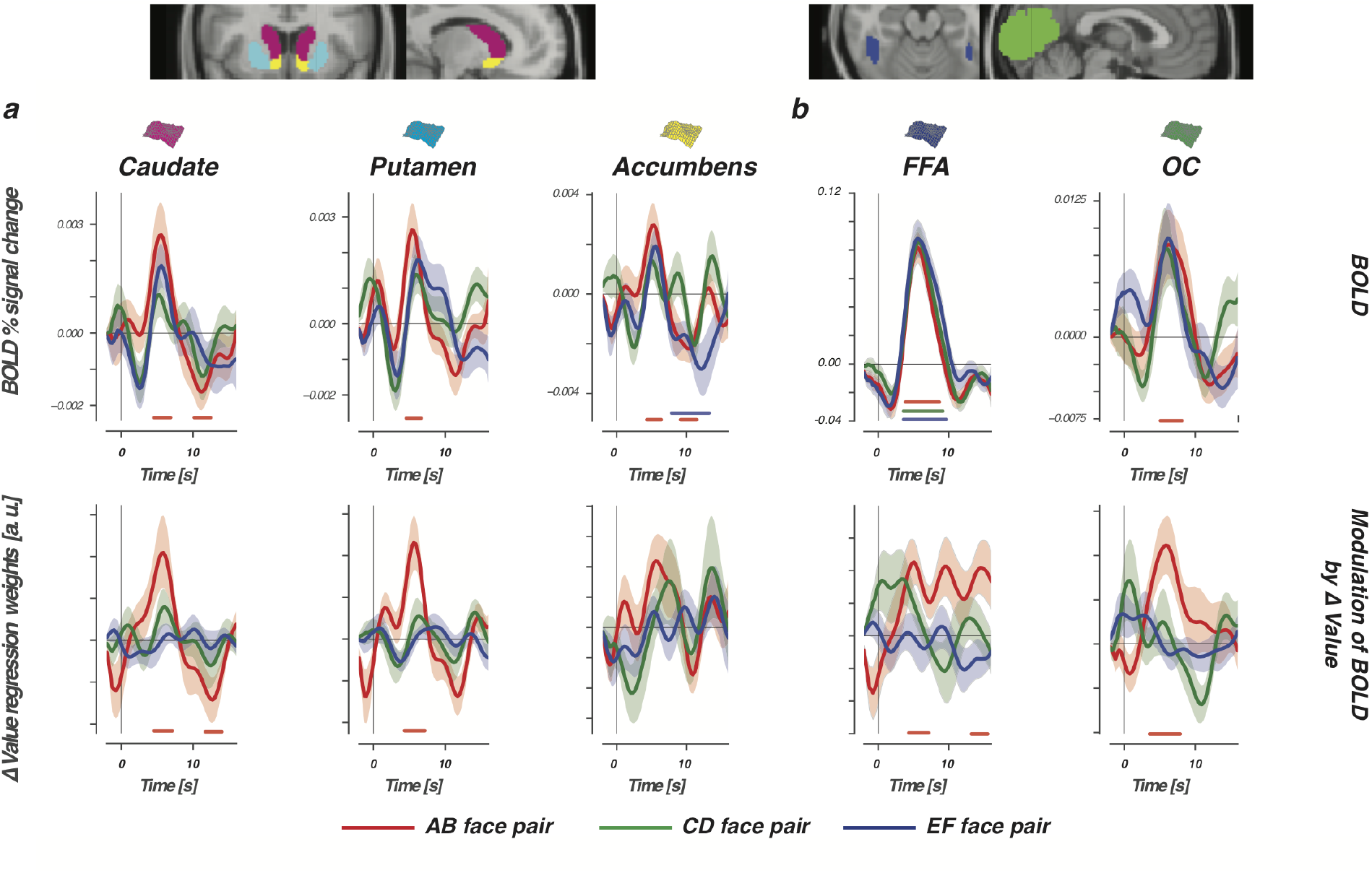
BOLD and the modulation of ∆ Value in the learning phase. Top row shows the BOLD signal time course, time-locked to presentations of AB (80:20, red lines), CD (70:30, green lines), and EF (60:40, blue lines) face pairs, for three striatal regions (**a**) and two perceptual regions (**b**). Bottom row displays differential modulation by value (∆Value = modulation *Q*_*chosen*_ – modulation *Q*_*unchosen*_). Horizontal lines show the interval in which modulation was significantly stronger for *Q*_*chosen*_. With the presentation of AB faces, BOLD responses in the dorsal striatum (caudate and putamen) and visual regions (FFA and OC) were modulated more by values of the chosen stimulus when compared to values of the unchosen stimulus. Differential AB value modulation was not significant in the ventral striatum (i.e., accumbens). Nor did we observe any differential value modulations with the presentation of the more uncertain CD and EF pairs. Confidence intervals were estimated using bootstrap analysis across participants (*n* = 1000), where the shaded region represents the standard error of the mean across participants (bootstrapped 68% confidence interval).

### Reward prediction errors in striatal and visual regions

Our findings so far described relationships between BOLD and value time-locked to the moment of stimulus presentation – i.e., when a choice is requested. Learning occurs when an outcome is different from what was expected. We therefore next focused on modulations of the BOLD response when participants received feedback. Learning modulations were explored by asking how trial-by-trial BOLD responses in perceptual and striatal regions relate to either signed (outcome was better or worse than expected) or unsigned (magnitude of expected violation) reward prediction errors (Fouragnan et al. 2018). Consistent with the literature, BOLD responses in all striatal regions were modulated by signed RPEs, with larger responses after positive RPEs or smaller responses after negative RPEs (Figure 5a bottom row). Activity in the accumbens (ventral striatum) was additionally tied to unsigned RPEs in the tail of the BOLD time-course, with larger violations (either positive or negative) tied to smaller dips. Consistently, estimated BOLD responses in both visual regions were modulated by the signed RPE, and once more mirrored the striatal modulations with stronger positive RPEs eliciting stronger BOLD responses (Figure 5b bottom row). FFA BOLD responses were additionally modulated by unsigned RPEs. However, in contrast to the relationship found between unsigned RPEs and the accumbens, the FFA modulation was positive and co-occurred with the modulation of the signed RPE. That is, bigger violations and more positive outcomes each elicited a stronger response in the FFA.

**Figure 5:**
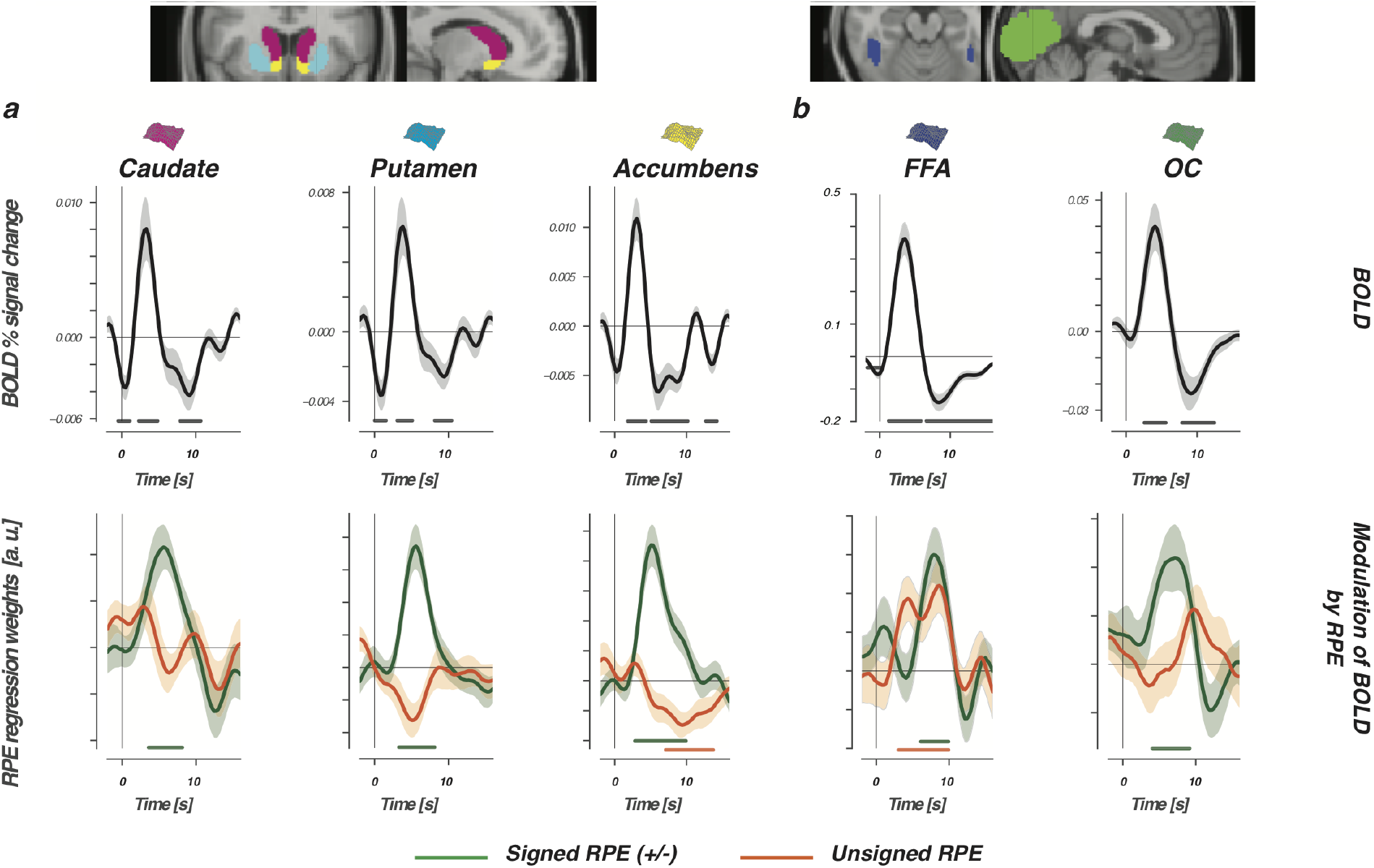
Reward prediction errors modulate BOLD in striatal and visual regions. The top row shows the FIR-estimated BOLD signal time-course, which was time-locked to the presentation of choice feedback and evaluated for three striatal regions (**a**) and two perceptual regions (**b**). Bottom row displays modulations of the estimated BOLD time-course by singed (green lines), or unsigned (orange lines) RPEs. The horizontal lines represent the interval in which signed or unsigned RPEs contributed significantly to the modulation of BOLD in the multiple regression. Note that both variables were always evaluated simultaneously in one GLM.

### Can past learning in visual regions support the prediction of future value-based decisions?

Stable value representations and reward prediction errors both modulated the activity of visual and striatal regions. These modulations in the striatum are described to bias future actions towards the most favored option (the dorsal striatum), or to predict future reward outcomes (the ventral striatum). To better understand the value and RPE modulations observed in visual regions, we next assessed the importance of these visual regions alongside the striatum in the correct classification (decoding) of future value-driven choice outcomes. Here, activity of prefrontal regions was added to the importance evaluation based on our previous work with this data in the transfer phase (Jahfari et al. 2018) (please see supplementary Figures 1&2 for the evaluation of these regions during learning).

In the transfer phase, participants had to make a value-driven choice based on what was learned before, i.e., during the learning phase. To specify the relevance of visual regions in the resolve of value-driven choice outcomes, in the transfer phase, a random forest (RF) classifier was used (Breiman 2001, 2004) (Please see Figure 2a-c for the procedure). The RF classifier was trained to predict the participant’s choice, on each trial, given trial-by-trial BOLD estimates from striatal, prefrontal, and visual regions. The RF classifier relies on an ensemble of decision trees as base learners, where the prediction of each trial outcome is obtained by a majority vote that combines the prediction of all decision trees (Figure 6a). To achieve controlled variation, each decision tree is trained on a random subset of the variables (i.e. subset of columns shown in Figure 2a), and a bootstrapped sample of data points (i.e. trials). Importantly, we ensured that the forest was not simply learning the proportion of optimal choices in the transfer phase by training all models on balanced draws from the training set with equal numbers of optimal and sub-optimal choices.

**Figure 6:**
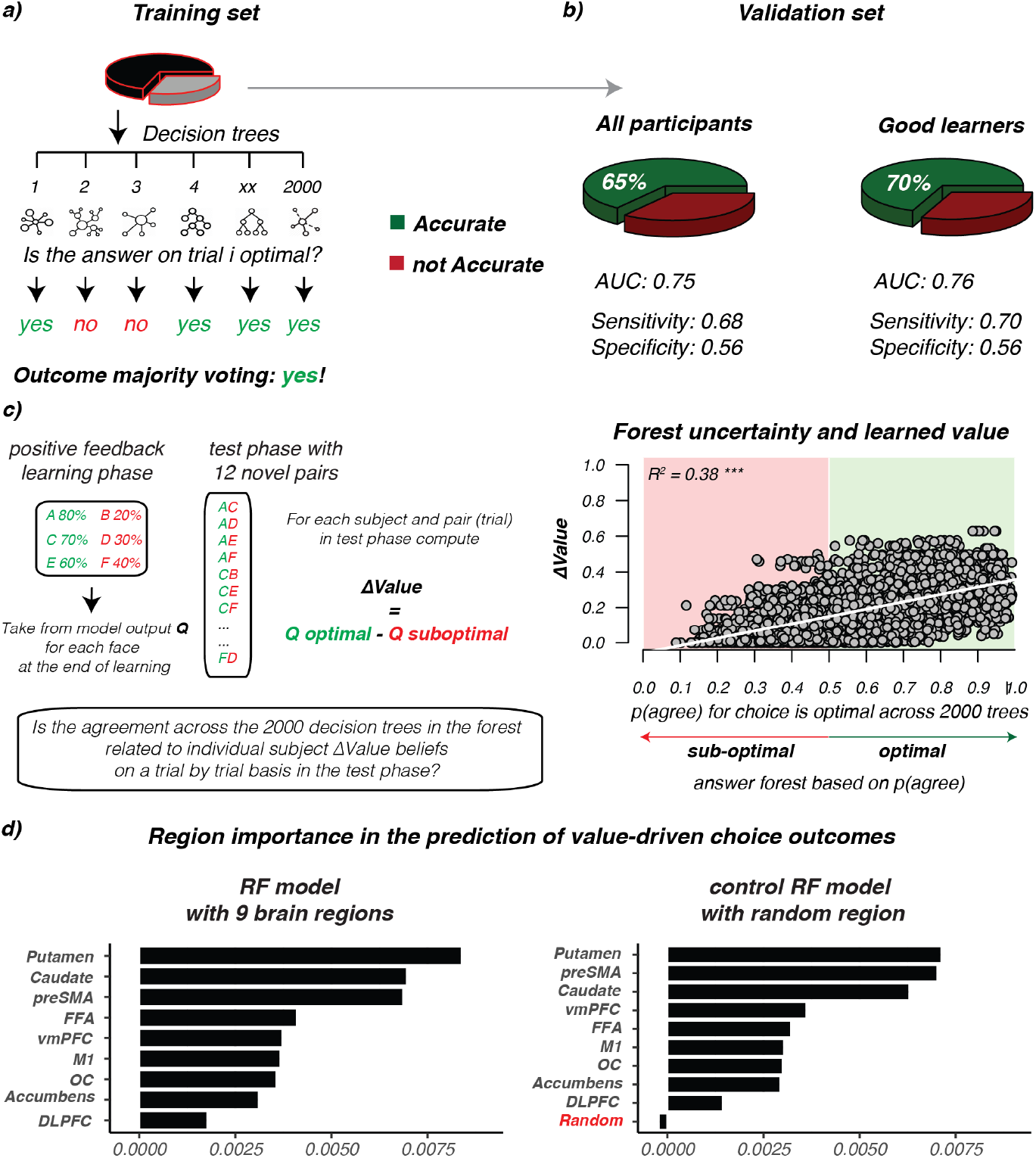
Random Forest performance and importance ranking. Prediction of value-driven choice outcomes in the transfer phase using trial-by-trial BOLD responses from striatal, perceptual, and prefrontal cortex regions. (**a**) Overview of the Random Forest approach where the training-set is used to predict choice outcomes for each trial by using the majority vote of 2000 different decision trees. Each tree is built using a different set, or sample, of trials and predictors from the training set. The forest is trained on a training set sampled from all participants (N=43), or only ‘the good learners’ (N=34). (**b**) Shows the classification, or decoding, accuracy (green) given the separate unseen validation sets, for all participants and good learners. (**c**) On the left, overview of the feedback scheme in the learning phase, and the new combination in transfer about which the RF is making an prediction with an illustration of how ∆Value is computed for each trial. ∆Value was computed for each trial in the transfer phase by using the end beliefs (*Q*) that participants had about each stimulus (A-to-F) at the end of the learning phase. On the right side, plotted relationship between forest uncertainty (i.e., proportion of agreement across 2000 trees), on each prediction/trial (x-axis) and ∆Value (y-axis) for the model with the highest accuracy (i.e., the good learners). Forest uncertainty is defined as the proportion of trees saying ‘yes! the choice on this trial was optimal/correct’. When this ratio is bellow 0.5 the forest will predict ‘no’ (sub-optimal/wrong choice), otherwise the prediction is ‘yes! the choice on this trial was optimal/correct’ (optimal). *R*^2^=adjusted *R*^2^. Note that, the same pattern was found for all participants (*R*^2^= 0.41^*∗∗∗*^, please see supplementary Figure 3). (**d**) Ranking of the ROI’s in their contribution to the predictive accuracy of the best performing model (i.e., good learners). Left, shows the original ranking. On the right, we evaluate ranking with all 9 original regions, but now add a control region that was sampled randomly from *N*(0, 1), and unrelated to the activity of any region, or ∆Value. Notice that the random variable has negative importance in the ranking, meaning that removing it improves model performance with 0.5%.

Evaluation of all participants resulted in a classification accuracy of 65% (*AUC* = 0.75) using the trial-by-trial BOLD estimates from the ROIs and increased to 70% with the evaluation of the good learners (*AUC* = 0.76; *N* = 34, criteria: accuracy > 60% across all three learning pairs). Hence, in 65 (all participants) or 70 (good learners) out of 100 trials the forest correctly classified whether participants would pick the option with the highest value (optimal choice) or not (sub-optimal choice) in the validation set. RF predictions were substantially lower when labels of the validation set were randomly shuffled (accuracy: all participants= 52%; good learners= 56%).

The improvement of accuracy with the evaluation of only the good learners is remarkable because the classifier was given less data to learn the correct labelling (fewer subjects/trials) and implied that the 2000 decision trees were picking up information related to the consistency of past learning. Further support for this important observation was found by asking how the uncertainty of each prediction (defined as the proportion of agreement in the predicted outcome among the 2000 trees for each trial) relates to the difference in value beliefs (∆Value) about the two options presented on each trial (computed using the end *Q*_*beliefs*_ of participants at the end of learning about face A-to-F), Figure 6c right side. As plotted in Figure 6c on the left, the uncertainty in predicting that a trial choice outcome is optimal – defined as the proportion of disagreement among the 2000 decision trees - decreased with larger belief differences in the assigned values (please see supplementary Figure 3 for the evaluation of all participants).

Besides providing insights into how BOLD responses in the transfer-phase contribute to predict value-driven choice outcomes (i.e., whether participants would choose the option with the highest value given past learning) the RF algorithm additionally outputs a hierarchy, thereby ranking the contribution of each region in the achieved classification accuracy. Figure 6d shows the ranking of all ROIs for good learners where the model had the highest predictive accuracy. First, regions in the dorsal striatum were most important, which aligned well with both the literature and the BOLD modulations we found by ∆Value and RPE during the learning phase. These regions were next followed by the preSMA. Evaluation of this region during the learning phase showed no modulations by ∆Value or RPE on BOLD (supplementary Figure 1&2). Nevertheless, this region is typically associated with choice difficulty/conflict and might be essential in the resolve of a choice when value differences are small. Remarkably, the third region in this hierarchy was the FFA. In a task where participants pick the most valued face based on past learning, this ranking of the FFA just above the vmPFC implies that the ∆Value and RPE modulations of BOLD observed during learning could function to strengthen the recognition of valuable features. With the evaluation of all participants – including some who were less good in learning – the ranking of both the FFA and vmPFC was much lower (please see supplementary Figure 3b), which might be caused by more noise across the group in learning.

Further insights in the role of perceptual regions came from the separate evaluation of RF for only the easiest (with ∆Value between the two choice options being large), or hardest (with small ∆Value) choices (supplementary Figure 6). Results showed that when ∆Value is large, or the choice is easy, RF predictions are best served by BOLD fluctuations in both dorsal and ventral striatum, followed by vmPFC, the preSMA and M1. With easy choices, regions involved with evidence accumulation (DLPFC), or perceptual processing (FFA and OC) rank last. More specifically, the processing of BOLD from OC even has a negative effect on RF accuracy, which means that running RF without OC will improve decoding. At the same time, with the evaluation of the most difficult choices - where participants decide between two very close in value positive (e.g., A or C) or negative (e.g., B or D) faces - we instead find perceptual regions to rank in the top. With difficult choices, where ∆Value is very small, the caudate is followed by the FFA and OC in serving RF predictions. We will return to the interpretation of these different rankings in the discussion.

Finally, we focused on two sets of control analysis. First, we evaluated RF accuracy and ranking with an additional random variable that was sampled from *N* (0, 1), and unrelated to the BOLD activity of any region, or ∆Value. Here, the added random control region ranks last with negative importance, meaning that removing it improves model performance with 0.5% (good learners) or 0.3% (all learners) points (right side Figure 6d, or supplementary Figure 3). Second, RF performance was evaluated with the removal of perceptual, striatal, or frontal regions. Despite the positive ranking of each region shown in Figure 6d (or supplementary Figure 3b), RF decoding was not affected by the removal of just one or two regions (supplementary Figure 5). However, accuracy is reduced when striatal (putamen, caudate, and accumbens), frontal (vmPFC, M1, DLPFC, and preSMA), or perceptual (FFA and OC) regions are evaluated in isolation. These alternative evaluations show that RF works best when trial-by-trial BOLD across multiple ‘learning’ brain regions is combined, but also that neither of the regions in isolation is crucial for the accuracy of predictions. Moreover, these control check highlight that when a variable is unrelated to learning, or single trial BOLD, ranking drops to last (as is to be expected) with counterproductive effects on RF accuracy.

## Discussion

This study provides novel insights into how reinforcements modulate visual activity and specifies its potential in the prediction of future value-driven choice outcomes. First, by focusing on how participants learn, we find BOLD in visual regions to change with trial-by-trial adaptations in value beliefs about the faces presented, and then to be subsequently scaled by the signed RPE after feedback. Next, the relevance of these observed value and feedback modulations was sought by exploring the prediction of future value-driven choice outcomes in a follow-up transfer phase where feedback was omitted. Our machine learning algorithm here shows a classification accuracy of 70% for participants who were efficient in learning by combining trial-by-trial BOLD estimates from perceptual, striatal, and prefrontal regions. The evaluation of region importance in these predictions ranked the FFA just after the dorsal striatum and the preSMA, thereby showing an important role for visual regions in the prediction of future value-driven choice outcomes in a phase where learning is established.

In a choice between two faces, BOLD responses in both the dorsal striatum and perceptual regions were affected more by values of the chosen face, relative to the unchosen face. Across three levels of uncertainty, we only observed the differential modulation of value on BOLD when belief representations were stable. This specificity aligns with neuronal responses to perceptual stimuli in the caudate tail (Kim et al. 2017), visual cortex (Shuler and Bear 2006; Weil et al. 2010; Cicmil et al. 2015), and imaging work across sensory modalities (Serences 2008; Serences and Saproo 2010; LimOdoherty2013; Pleger et al. 2009; Kahnt et al. 2011; Vickery et al. 2011; FitzGerald et al. 2013; Kaskan et al. 2016), where it fuels theories in which the learning of stable reward expectations can develop to modulate, or sharpen, the representation of sensory information critical for perceptual decision making (Roelfsema et al. 2010; Kahnt et al. 2011; Cicmil et al. 2015).

After a choice was made, feedback modulations of signed (‘valence’) and unsigned (‘surprise’) RPEs (Fouragnan et al. 2018) were evaluated on BOLD responses, by using an orthogonal design where the unsigned and signed RPE compete to explain BOLD variances. Both visual and striatal regions respond to prediction errors (Den Ouden et al. 2012). In the striatum both valence and surprise are thought to optimize future action selection in the dorsal striatum, or the prediction of future rewards in the ventral striatum. In perceptual regions, a mismatch between the expected and received outcome is often explained as surprise where a boost in attention or salience changes the representation of an image without a representation of value per se. We found positive modulatory effects of signed RPEs in all striatal regions, as well as, in the FFA and OC. Concurrently, modulations of unsigned RPEs were only observed in the accumbens (ventral striatum) and FFA, where notably the direction of modulation was reversed. We speculate that this contrast arises from the differential role of the regions. In the FFA, specialized and dedicated information processing is essential to quickly recognize valuable face features. Complementary boosts of surprise and valence here could prioritize attention towards the most rewarding face feature to strengthen the reward association in memory, or help speed up future recognition (Gottlieb 2012; Gottlieb et al. 2014; Störmer et al. 2014). In the accumbens, boosted effects of positive valence on BOLD were dampened by larger mismatches. Large mismatches in what was expected are rare in stable environments. We therefore reason that in the accumbens the contrast between valence and surprise could function as a scale to refine learning, eventually leading to more reliable predictions of future rewards.

Whereas BOLD in the ventral striatum was shaped by both signed and unsigned RPEs, the dorsal striatum was sensitive to differential value up-to a choice and signed RPEs with the presentation of feedback (Kaskan et al. 2016; Lak et al. 2016, 2017; McCoy et al. 2018; Van Slooten et al. 2018). The concurrent modulation of differential value in the primary motor cortex (please see M1 in supplementary Figure 1) associates the dorsal striatum with the integration of sensory information (Ding and Gold 2010; Yamamoto et al. 2012; Hikosaka et al. 2013; Kim et al. 2017), where increased visual cortex BOLD responses to faces with the highest value could potentially help bias the outcome of a value-driven choice.

We explored this line of reasoning with the prediction of value-driven choice outcomes in a follow-up transfer phase after leaning. In recent years, machine learning approaches have become increasingly important in neuroscience (Naselaris et al. 2011; Hassabis et al. 2017; Hebart and Baker 2018; Snoek et al. 2019), where the ease of interpretation has often motivated a choice for linear methods above non-linear methods (Naselaris et al. 2011; Kriegeskorte and Douglas 2018). Despite the latter being less constrained and able to reach a better classification accuracy by capturing non-arbitrary, or unexpected relationships (King et al. 2018). Value-driven choices after a phase of initial learning are influenced by the consistency of past learning, memory updating, and attention. All of these processes are affected by both linear and non-linear neurotransmitter modulations (Aston-Jones and Cohen 2005; Yu and Dayan 2005; Cools and D’Esposito 2011; Beste et al. 2018). Our RF approach was unconstrained by linearity with classification accuracies well above chance and improved with the evaluation of only the good learners; despite substantial decreases in data given to the algorithm to learn the correct labelling. Critically, we additionally found that the uncertainty of trial-by-trial predictions made by RF is tied to the differentiability of value beliefs – an index that we could compute for the novel pair combination in the transfer phase by using the value (*Q*) beliefs that participants had about each face at the end of learning. These results showcase how trial-by-trial BOLD fluctuations in striatal, prefrontal, and sensory regions can be combined by machine learning, or decoding, algorithms to reliably predict the outcome of a value-driven choice. Where we refine the interpretation of non-linear predictions by combining the RF output with cognitive computational modelling. With this combination we essentially show how the uncertainty of RF predictions is tied to value beliefs acquired with learning in the past.

An important evaluation intended with our machine learning approach was the ranking of regions by their contribution to the predictive (decoding) accuracy in the transfer phase. After the observed modulations of BOLD in the learning phase this explorative analysis sought the relevance of learning-BOLD relationships in the resolve of future choices. Here, the ranking made by RF first identified signals from the dorsal striatum (putamen and caudate) as most important followed by the preSMA, and then most notably, visual regions. That is, when the quality of leaning was high across participants, FFA ranked just above traditional regions such as the vmPFC and the accumbens (O’Doherty et al. 2003, 2017; Hare et al. 2011; Niv et al. 2012; Klein et al. 2017). Notably, FFA was replaced by OC in ranking with the evaluation of all participants (please see supplementary Figure 3b). This difference could occur because the quality of learning was more variable across all participants, or because RF predictions based on the heterogeneous data from all participants were less accurate. In general, the shift in ranking implies that when learning is less consistent choice outcomes are better predicted by fluctuations in OC - perhaps with the identification of rewarding low-level features. With better or more consistent learning, however, participants should increasingly rely on memory and specialized visual areas. Thus, search for specific face features associated with high value by recruiting the FFA in the visual ventral stream. Consistent with this reasoning recent neuronal recordings show rapid visual processing of category-specific value cues in the ventral visual stream. These specific value cues are only seen for well-learned reward categories, and critically, precede the processing of value in prefrontal cortex (Sasikumar et al. 2018).

Additionally, in the learning phase both OC and the FFA were modulated more by values of the (to be) chosen stimulus when belief representations were stable and distinct - i.e., we only observed differential *Q*-value modulations for the most reliable and easy to learn AB pair. This combined with the RPE modulations found in the same regions suggests an effect of value and learning on perceptual regions that is both specialized (FFA) and global (OC). Note however that this possibility must be studied further with designs that can zoom in on specificity with the separation of different perceptual dimensions (e.g., houses vs faces). Our transfer phase resluts imply a differential role for the specialized FFA, and the more low-level general OC, with the comparison of good vs all learners. Tasked with predicting the outcome of future value-driven choices RF rankings showed a specialized and prominent FFA role for good/efficient learners whereas OC was more important with the evaluation of all participants (where learning was less consistent or noisier across participants). Recent work on the interplay between learning and attention suggests a bi-directional relationship between learning and attention: we learn what to attend from feedback, and in turn, use selective attention to constrain learning towards relevant value dimensions (Leong et al. 2017; Rusch et al. 2017). In our study, better learning helps a more refined identification of rewarding features in a face, which we interpret as a narrower focus of selective attention in the FFA during learning (Niv et al. 2015). With past learning being more noisy, or less established, extraction of relevant features is less straightforward with attention being more spread to both specialized and global regions. Additionally, we observed both FFA and OC to only rank in the top (just after the caudate) when ∆Value was very small (supplementary Figure 6). With easy choices this effect was reversed where processing of OC BOLD even declined the RF accuracy. This contrast suggests, that especially when the options to choose from are just too similair in value (i.e., think of the options A:C, or B:D), past learning in perceptual regions could serve the striatum with a selective boost to highlight the most rewarding face features. In contrast, when the distinction is easy and clear-cut, choices depend far more on inputs from the ventral striatum and vmPFC.

We note that although BOLD fluctuations in the preSMA ranked second in the prediction of value-driven choice outcomes, no reliable modulations of BOLD were observed by either differential value or RPEs in the learning phase. The preSMA is densely connected to the dorsal striatum and consistently associated with action-reward learning (Jocham et al. 2016), or choice difficulty (Shenhav et al. 2014). The lack of associations in this study might result from our noisier estimates of the BOLD response that is typical for regions in the prefrontal cortex (Pircalabelu et al. 2015; Bhandari et al. 2018), the anatomical masks selected, or smaller variability across trials in the learning phase (i.e., 3 pairs in learning-phase vs 15 pairs in transfer-phase). Nevertheless, the importance indicated by RF, combined with our previous analysis of this transfer phase data (Jahfari et al. 2018), implies an important role for the preSMA in the resolve of value-driven choices in concert with the striatum. More research with optimized sequences to estimate BOLD in PFC is required to clarify the link between learning and transfer.

To summarize, we find an important role for perceptual regions in the prediction of future value-driven choice outcomes, which coincides with the sensitivity of BOLD in visual regions to differential value and signed feedback. These findings imply visual regions to learn prioritize high value features with the integration of feedback, to support and fasten, optimal response selection via the dorsal striatum in future encounters.

## Acknowledgements

This work was supported by an ABC Talent grant to SJ from the University of Amsterdam, an ERC grant ERC-2012-AdG-323413 to JT, and NWO-CAS grant 012.200.012 to TK.

## Author contribution

SJ and TK developed the questions and analysis plan for the re-analysis. SJ and TK contributed novel methods and analyzed the data. SJ wrote the first draft of the MS with edits from TK. JT commented on the final draft.

## Data availability

The code and preprocessed files for behavioral and decoding analyses can be download from: https://github.com/sarajahfari/Pearl3T.git, and fMRI preprocessing and deconvolution analysis code are available at https://github.com/tknapen/pearl_3T. The raw data can be downloaded from openneuro.org in BIDS after acceptance of this MS.

