## supplementary figure for "Learning in visual regions as support for the bias in future value-driven choice"

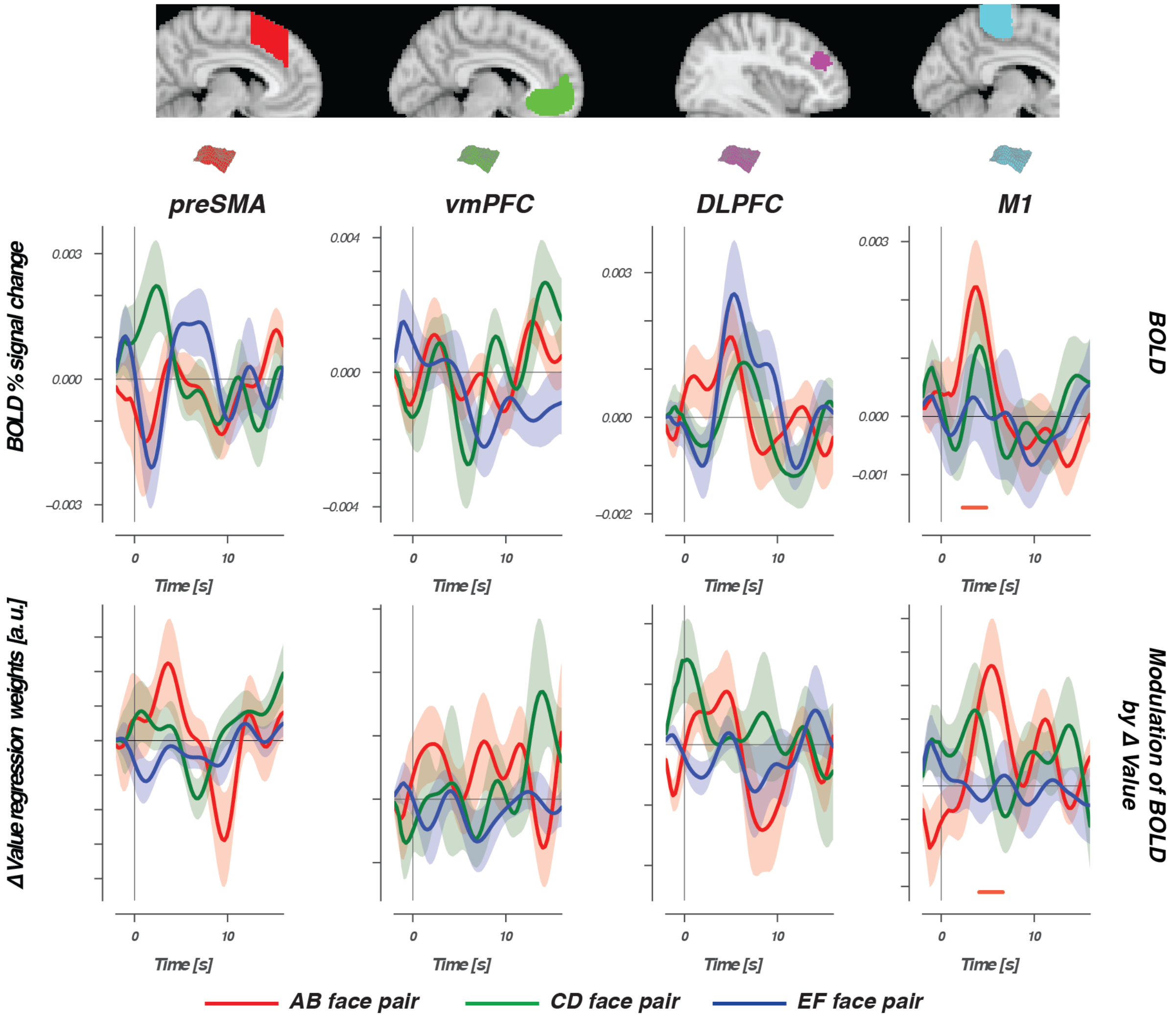


Figure 1: **BOLD and the modulation of ΔValue in the learning phase for cortical regions.** The top row shows the FIR-estimated BOLD signal time-course, time-locked to the presentation of AB (red lines), CD (green lines), and EF (blue lines) face pairs for the additional cortical regions that were evaluated with RF. The bottom row displays the differential modulation by value (ΔValue = modulation Q chosen − modulation Q unchosen). The horizontal lines show the interval in which the modulation was significantly stronger for Q chosen. With the presentation of AB faces, only the BOLD responses in M1 was modulated more by values of the chosen stimulus when com-pared to values of the unchosen stimulus. Confidence intervals were estimated using bootstrap analysis across participants (n=1000), where the shaded region represents the standard error of the mean across participants (i.e. bootstrapped 68% confidence interval).

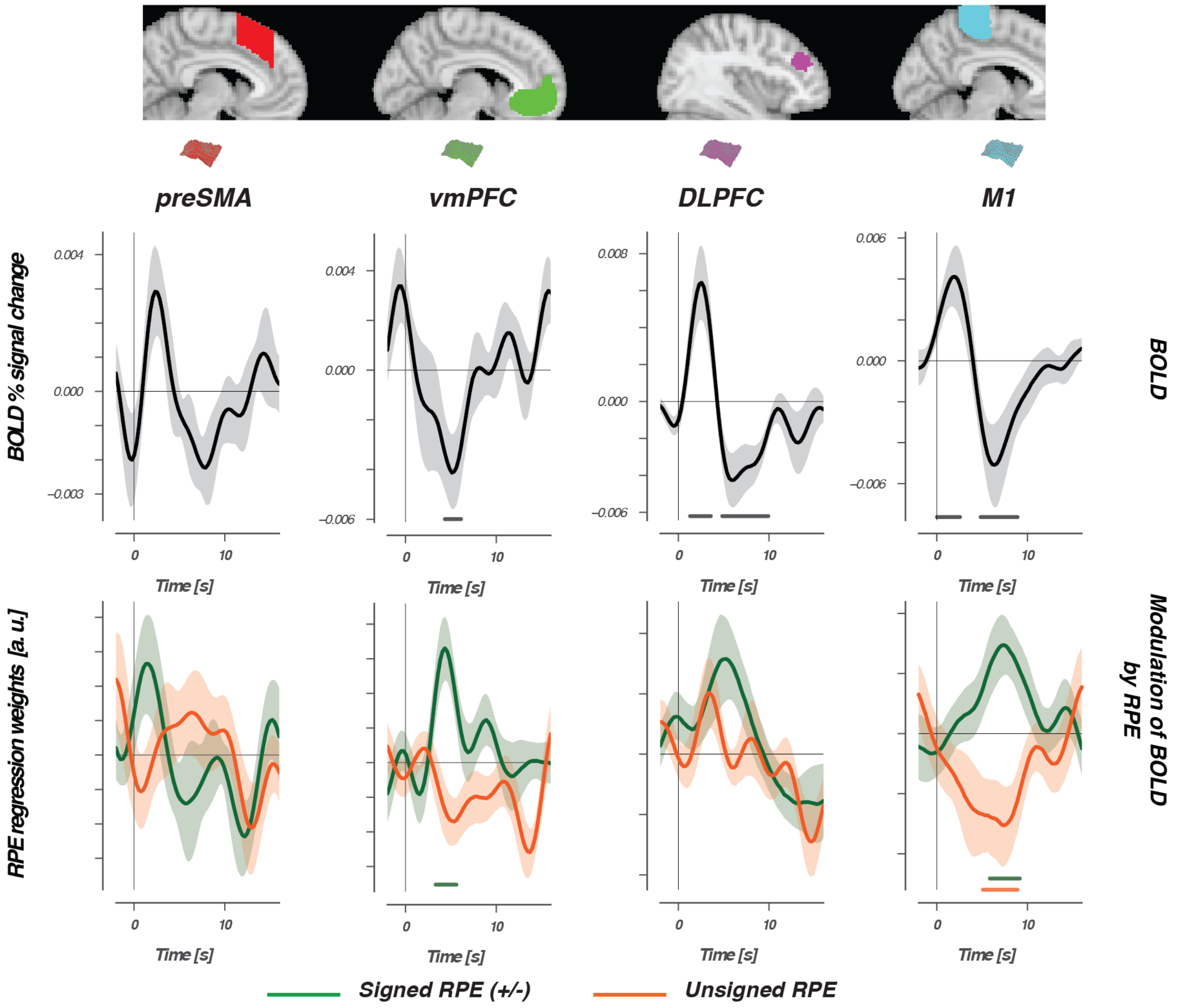


Figure 2: **Reward prediction errors and BOLD during the learning phase in cortical regions.** The top row shows the FIR-estimated BOLD signal time-course, which was time-locked to the presenta-tion of choice feedback, shown for the additional cortical regions evaluated with RF. The bottom row displays modulations of the estimated BOLD time-course by singed (green lines), or unsigned (orange lines) RPEs. The horizontal lines represent the interval in which singed or unsigned RPEs contributed significantly to the modulation of BOLD in the multiple regression. Note that, both variables were always evaluated simultaneously in one GLM.

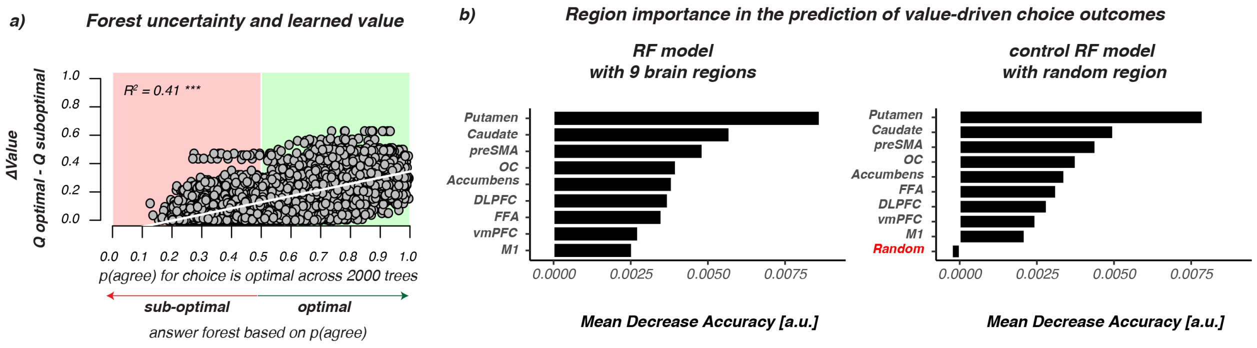


Figure 3: **Uncertainty and Ranking of RF evaluated for all participants.** **a**) Plotted relationship between the uncertainty of the forest in each prediction (x-axis) and delta value (y-axis) for all participants. ΔValue was computed for each transfer-phase trial using the end ($Q$) beliefs that participants had about each face at the end of the learning phase. Forest uncertainty is defined as the proportion of trees saying 'yes! the choice on this trial was optimal'. When this ratio is bellow 0.5 the forest will predict 'no' (sub-optimal), otherwise the prediction is 'yes! the choice on this trial was optimal' (optimal). $R^2$=adjusted $R^2$. **b**) Plotted ranking of the ROI's in their contribution to the predictive accuracy of RF evaluated with all participants. Left, the original ranking for all participants. Right, the ranking with the 9 original plus a control region that was sampled randomly from $\mathcal{N}(0, 1)$, and unrelated to the activity of any region or ΔValue. As with the good learners the random variable has negative importance in the ranking, meaning that removing it improves model performance with 0.3% points.

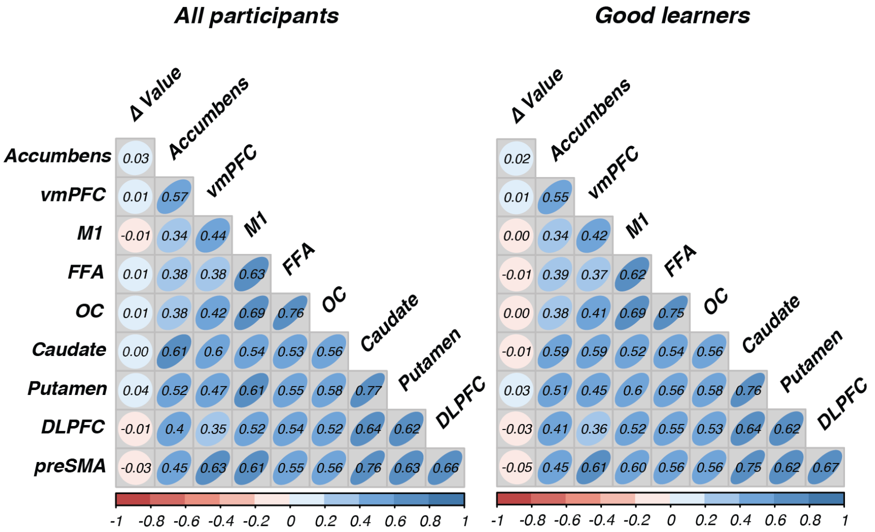


Figure 4: **Correlation Matrix for single trial BOLD across regions, or with ΔValue**. The left panel evaluates all participants (N=43), with the good learners (N=34) on the right. Centroids combine shape and strength of the relationship with the number on top of each box being the correlation value. It stands out that across all brain regions trial-by-trial BOLD is correlated, where we see the strongest relationships between regions of the dorsal striatum (i.e., Caudate and Putamen), visual cortex (i.e., OC and FFA), or the PFC and the dorsal striatum (i.e., Caudate and preSMA or DLPFC). These observations concur with our connectivity evaluations of this data showing effective connectivity inputs from the preSMA and DLPFC into the dorsal striatum, and earlier work reporting functional and effective connectivity between the FFA and striatum. Also note the lack of relationship between ΔValue single trial BOLD from any region. This contrasts the relationship between ΔValue and RF predictions, where the latter combines trial-by-trial information across regions to predict a choice.

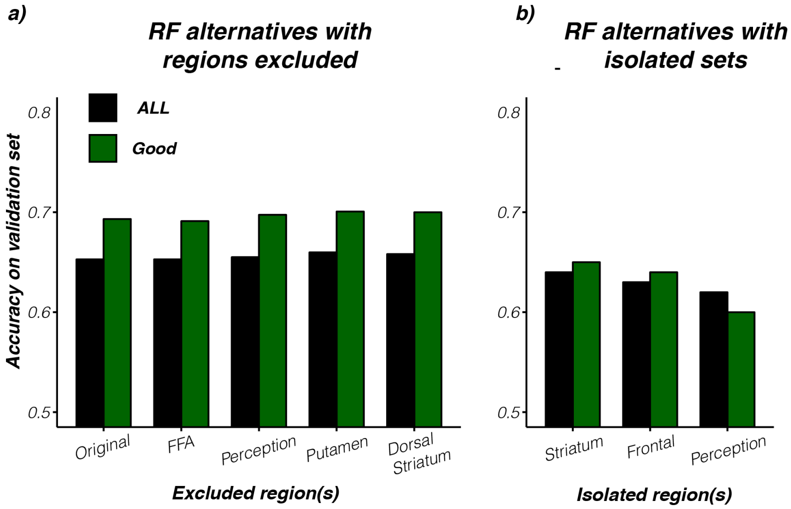


Figure 5: **Alternative evaluations with RF**. **a**) RF classification accuracy is not reduced with the removal of perceptual, or striatal regions when compared to the original model with 9 ROI's. Perception = exclusion of OC and FFA, Dorsal Striatum = exlusion of the Putamen and Caudate. **b**) However, accuracy is lower when striatal (Putamen, Caudatae, and Accumbens), frontal (vmPFC, M1, DLPFC, and preSMA), or perceptual (FFA and OC) regions are evaluated in isolation. These alternative evaluations show that RF works best when trial-by-trial BOLD across multiple brain regions is combined.

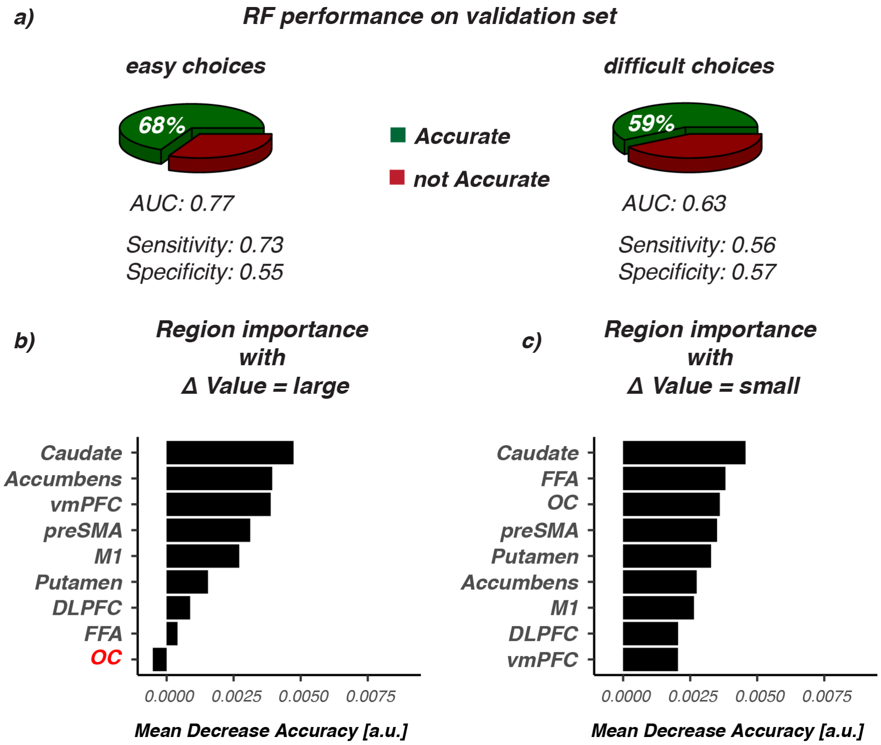


Figure 6: **Random Forest performance and ranking split for easy and difficult choice trials** . For this analysis we created an easy choice set - with ΔValue between the two choice options being large given participants beliefs at the end of learning - a medium choice set (not evaluated), and a very difficult choice set where ΔValue was very small. RF was run separately for the easy and hard set with balanced numbers of correct and incorrect choices, where we again set aside 1/3 of each set for validation. Because the trial-by-trial BOLD data is cut in three this evaluation included all participants (N=43). **a**) Shows the classification or decoding accuracy (green) given the separate unseen validation sets for 'easy' value-driven choices where ΔValue is large (e.g., a choice between A and D, or C and B), or 'hard' choices with small ΔValues (e.g., between A and C, or B and D). **b**) Plotted ranking of the ROIs from the RF model for easy, or hard choices (**c**). Notice that with easy choices, OC has a negative ranking. Removing OC here improves RF decoding.
